# Genotypic identification of polyclonal plasma cells in plasma cell dyscrasias shows an aberrant single-cell phenotype with clinical implications

**DOI:** 10.1101/2024.05.26.595470

**Authors:** Matteo Claudio Da Vià, Francesca Lazzaroni, Antonio Matera, Alessio Marella, Akhiro Maeda, Claudio De Magistris, Loredana Pettine, Sonia Fabris, Stefania Pioggia, Alfredo Marchetti, Marzia Barbieri, Silvia Lonati, Alessandra Cattaneo, Marta Tornese, Margherita Scopetti, Nayyer Latifinavid, Giancarlo Castellano, Federica Torricelli, Antonino Neri, Cathelijne Fokkema, Tom Coupedo, Marta Lionetti, Francesco Passamonti, Niccolò Bolli

## Abstract

Multiple Myeloma (MM) is driven by clonal plasma cell (PC)-intrinsic factors and changes in the tumorigenic microenvironment (TME). To investigate if residual polyclonal PCs (pPCs) are disrupted, single-cell (sc) RNAseq and sc B-cell receptor analysis were applied in a cohort of 46 samples with PC dyscrasias and 18 healthy donors (HDs). Out of n=*213,074* CD138^pos^ PCs, 42,717 were genotypically identified as pPCs. Compared to HDs, we detected quantitative and qualitative differences in pPCs of patients showing immunoparesis, where we showed a pro-inflammatory status, driven by specific cellular interactions with TME. Finally, we derived a “hPC signature” that, once inferred in the CoMMpass dataset, was predictive of PFS and OS. Our findings show that genotypic, single-cell identification of pPCs in PC dyscrasias has relevant pathogenic and clinical implications.

## INTRODUCTION

Multiple myeloma (MM) is a plasma cell (PC) neoplasm whose origin is in the germinal center of a secondary follicle, in a B-cell undergoing class-switch recombination and somatic hypermutation upon antigen encounter^1,2^. Initiating somatic events at the genomic level are thought to be translocations of recurrent oncogenes to the immunoglobulin heavy chain (IGH) locus, or trisomies of several odd-numbered chromosomes^1^. The genomically aberrant post-germinal center B-cell will still migrate to the bone marrow (BM), differentiate into a (monoclonal) antibody producing PC, and initiate a monoclonal expansion^2^. These early events in myeloma development are not recognized at the clinical level. However, an initial BM expansion of clonal PCs can be easily identified by detecting a monoclonal protein (MP) with peripheral blood (PB) serum protein electrophoresis. Asymptomatic PC expansions can be categorized as monoclonal gammopathy of undetermined significance (MGUS) or smoldering multiple myeloma (SMM) based on the BM monoclonal PC percentage^3–5^. MGUS and SMM are thought to universally precede development of active MM, even when not recognized. In turn, active MM is a clinically aggressive neoplasm that has caused end-organ damage or shows surrogate signs of high disease burden with imminent risk of end-organ damage^5^, and requires treatment to prolong the patient’s life. Evolution from MGUS to SMM and MM is paralleled by the acquisition of secondary somatic genomic events, such as gene mutations, further translocations and copy-number abnormalities (CNAs)^6–8^. However, the serial study of MM samples evolved from SMM shows that some cases do not display genomic changes^9,10^, so that not all clinical progressions can be explained by genomics alone. Indeed, the tumor microenvironment (TME) plays a clear pathogenic role in MM development^11,12^. Clonal PCs have been shown to interact with immune and non-immune microenvironment cells through aberrant signaling, receiving in turn pro-survival and anti—apoptotic paracrine stimuli^13–16^. Furthermore, surveillance from immune TME cells has the potential to prevent progression of asymptomatic stages of clonal PC expansion^17^, and is disrupted along its progression^11,15,18,19^. Indeed, animal models show how loss of immune surveillance is causal to MM development from precursor stages^20^. Strategies to restore anti-MM immune function are a promising treatment approach in relapsed-refractory MM (RRMM) cases, where high-risk genomic lesions are enriched and response to conventional treatments is poor^21–23^. In the era of novel immunotherapies, having a fit immune system correlates with the quality of response, while T-cell exhaustion predicts poor response to T-cell engagers and chimeric antigen receptor T cells^24–28^. Immunoparesis, defined as reduced levels of one or more non-clonal immunoglobulin classes, is frequent in PC dyscrasias. Its prevalence increases from MGUS to SMM and MM. In asymptomatic conditions, its presence is prognostic for evolution^29–31^. In MM patients, immunoparesis can explain the increased susceptibility to infection and is associated with an overall^32^ worse prognosis^31^. Clearly, immunoparesis can only be explained by disruption of polyclonal PCs. Indeed, a reduction in the number of phenotypically-defined polyclonal PCs has been described along progression of PC dyscrasias. Less is known about the functional properties of polyclonal PCs (pPCs) in various stages of PC dyscrasias. Single cell RNA sequencing (scRNAseq) has the potential to dissect the clonal and polyclonal PC populations, and initial studies have found a different transcriptomic signature of the two populations^33–35^. However, our studies and others ^34,35^ have shown that polyclonal PCs do not always clearly separate from clonal PCs in scRNAseq Uniform Manifold Approximation and Projection (UMAP) plots, suggesting that a pure transcriptomic approach fails to accurately identify pPCs. Newer technologies allowing single-cell B-cell receptor (BCR) genotyping along with transcriptome sequencing of the same single-cell are better suited to unambiguously identify pPCs. In this paper, we developed an algorithm to unambiguously identify polyclonal PCs at the single-cell genotypic level. This allowed us to study their functional properties in MGUS, SMM and MM stage as well as in comparison with normal PCs from healthy donors (HD), showing they have profoundly deranged functions whose identification is relevant for understanding disease pathogenesis and has prognostic implications.

## RESULTS

### Landscape of polyclonal plasma cells across stages of PC dyscrasias

We analyzed CD138-purified samples from 7 MGUS, 16 SMM and 23 MM patients. CD138+ samples from HD were sourced locally (n = *1*), from collaborators (n = *3*), and from publicly available databases (*n* = 14)^34,36^. For patients with PC dyscrasias (**Table S1)**, we developed a workflow (**Figure 1A**) to leverage single-cell data to genotypically identify clonal vs polyclonal PCs. The polyclonal nature of the latter group was confirmed by the lack of identity and hierarchical relationship of the heavy and light chain BCR sequence with the BCR sequence of clonal PCs of the same patient (**Table S2**). PCs sharing the same light chain, or substantial identity with the clonal sequence were excluded from analysis (**STAR Methods, Table S2**). Copy-number abnormalities (CNAs) analysis through InferCNV^37^ confirmed that pPCs of each individual sample did not share any of the CNAs observed in their cPCs counterparts (**Figure S1,S4**). After filtering cells using standard quality controls, a total of n= *213,074* PCs (n=*32,704* from HDs, n= *15,882* from MGUS, n= *55,409* from SMM and n= *109,079* from MM) (**Figure 1B**) were used for further analysis. Using this approach, n= *170,357* cells (79,95%) were genotyped as clonal (cPCs) and n= *42,717* (20.04%) as polyclonal (pPCs). Overall, the frequency of pPCs was higher for MGUS and decreased in SMM and MM samples. The number of pPCs was n= *3,103* (19.53%) in MGUS, n= *3,291* (5.93%) in SMM, and n= *3,619* (3.31%) in MM (**Figure 1B-C)**. In the sample space, pPCs from different individuals tended to cluster together. However, there was a considerable admixture of pPCs and cPCs in the sample space of each patient (**Figure 1D**), indicating that transcriptomics alone is likely insufficient to accurately dissect every pPC from cPCs. The UMAP distribution of cells by disease stage also lacked a clear-cut picture, even though MM samples tended to differentiate the most from the asymptomatic ones (**Figure 1E**). We next looked at canonical PC marker genes. Our analysis confirmed previous observations, such as the frequent loss of *CD27*^34,38,39^ in cPCs, and their uniform expression of *CD38*, *CD138* and *BCMA (TNFRSF17)*. As expected, oncogenes such as *NSD2* and *CCND1/2,* and *MAF* were only expressed in cPCs. Interestingly, we found *GPRC5D* expression to be on average lower in pPCs than in cPCs from the same patients (**Figure S5A-B**) which might have clinical implications for the risk of infections in anti-GPRC5D T-cell redirecting immunotherapies versus anti-BCMA treatment^25^ (**Figure S5C-D**). Of note, pPCs frequently expressed markers usually attributed to cPCs such as *IGTB7*, *CCND3*, and this was true for both pPCs from patients and PCs from HDs (**Figure 1F**). In summary, these results highlight the importance of clonotypically identifying cPCs from pPCs in patients with PC dyscrasias. While canonical marker genes show expected expression patterns, whole-transcriptome analysis shows that pPCs have a transcriptomic profile that is not clear-cut different from cPCs.

**Figure 1.**
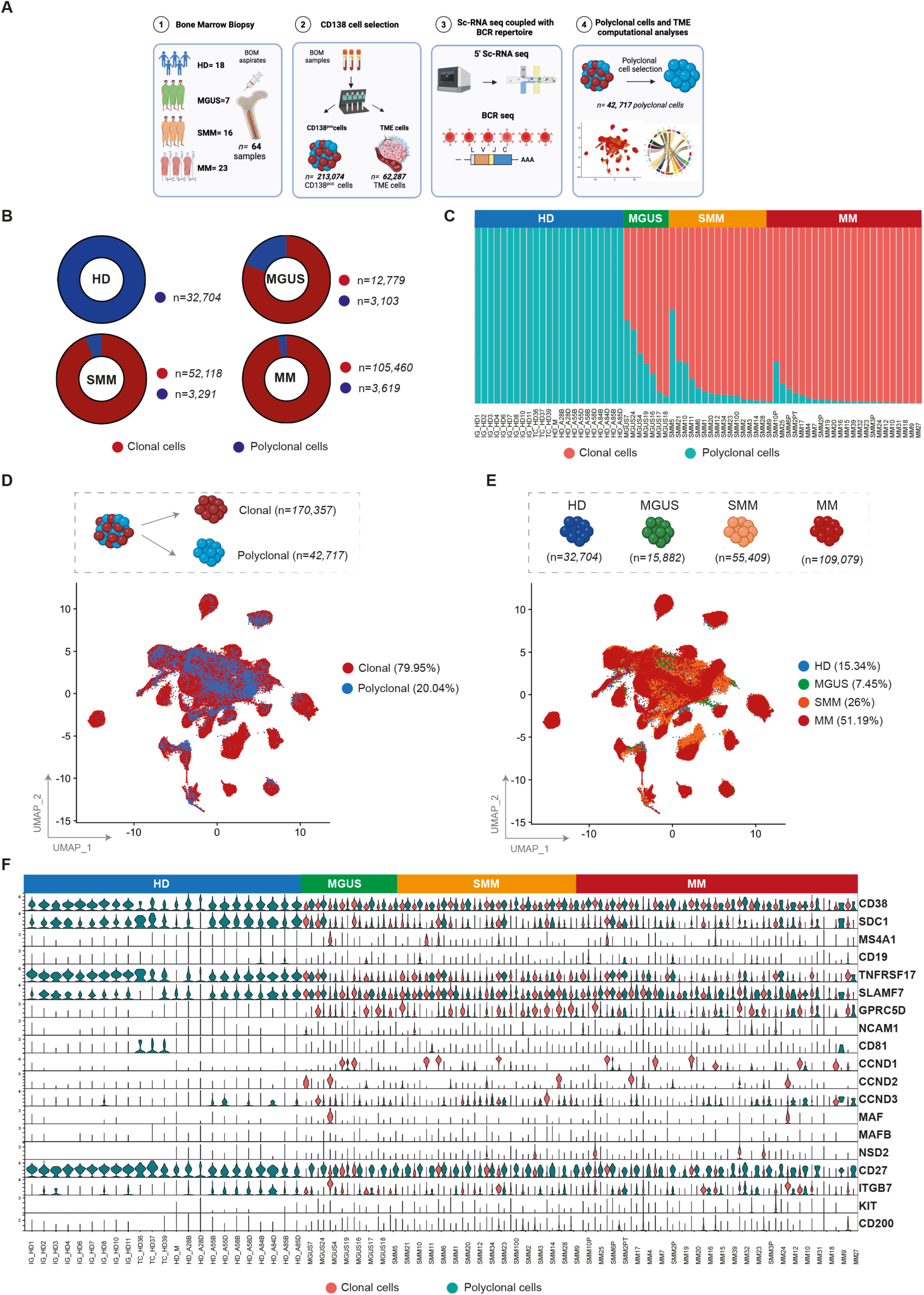
Clonal and polyclonal cells identification in patients affected by plasma cell dyscrasias and healthy donors (HD). **(A)** Schematic overview of cohort of samples and workflow of the study. **(B)** Donut charts representing the distribution of clonal and polyclonal cells in plasma cell dyscrasias and healthy donors based on clinical stratification (HD: blue; MGUS: green; SMM: orange; MM: red). Numerical values are reported in the legend for category. Color legend of clonal and polyclonal cells as in (B). **(C)** The proportions of clonal cells (red) and polyclonal cells (blue) in each sample. **(D)** UMAP plot show all clonal (red) and polyclonal (blue) cells, with numerical values. **(E)** UMAP representation of as per **(D),** colors indicate clinical stratification of samples. Numerical values and percentages are reported in the legend for each category. **(F)** Violin plots showing distribution of expression of genes commonly upregulated in patients plasma cell dyscrasias, along with annotations of the clinical classification (top). Bar above violin plots summarize clinical type assignments. Color legend as in (**B** and **C**).

### Polyclonal PCs from patients show deregulated expression profiles and an inflammatory phenotype

To further delineate the phenotype of n= *42,717* pPCs, we started the analysis by determining transcriptomic heterogeneity of pPCs in our cohort of samples, after filtering cells using standard quality controls (**Figure S6A-B**). Re-plotting only pPCs from patients and PCs from HDs, we observed a slight divergence of the two groups, but overall, there was a considerable overlap of cells from all disease stages (**Figure 2A**). After creating UMAP plots by clusters and by patients (**Figure S7A-B**), we next asked what genes are differentially expressed in the different clinical categories. Interestingly, pPCs from patients showed a similar pattern, different from HDs (**Figure 2B**). The latter group showed overexpression of genes such as *DUSP1*, *SELK*, *FKBP2* and *GNB2L1*, mainly involved in protein folding and protection against oxidative stress. Conversely, in pPCs from patients we found upregulation of genes usually associated with inflammation and/or stress conditions, such as the autophagy receptor *SQSTM1*, the growth inhibitor *GADD45B*, and the *VIM* gene, often upregulated during epithelial to mesenchymal transition (**Figure 2B)**. When looking at sets of genes implied in MM pathogenesis, again canonical PC marker genes were not different between pPCs in patients and HDs. We recognized exclusively genes in pPCS from patients that are not expressed in HDs such as *CTSB*, *CTSD*, *OPTN* involved in autophagy. Even more interestingly, we found that only PCs from SMM and MM upregulate the surface integrin *ICAM1*, implying that interaction with the TME changes for pPCs in more advanced stages (**Figure 2C**). We performed a differential expression (DE) analysis to look at pathway analyses in all possible 1:1 contrast combination (Wilcoxon rank sum test, with *padjust* Benjamini-Hochberg correction). Compared to pPCs from HD there was an upregulation of inflammatory pathways (hallmark TNF α and IFN α response, hallmark oxidative phosphorylation) in pPCs from pathological samples, as well as comparing pPCs across disease stages to each other, with the most clear-cut differences between MGUS and SMM/MM (**Figure 3A, Figure S8A-C)**. In summary, we showed a *climax* of increasing inflammatory status from HD to MM (**Figure 3B, Figure S8A-C)**.

**Figure 2.**
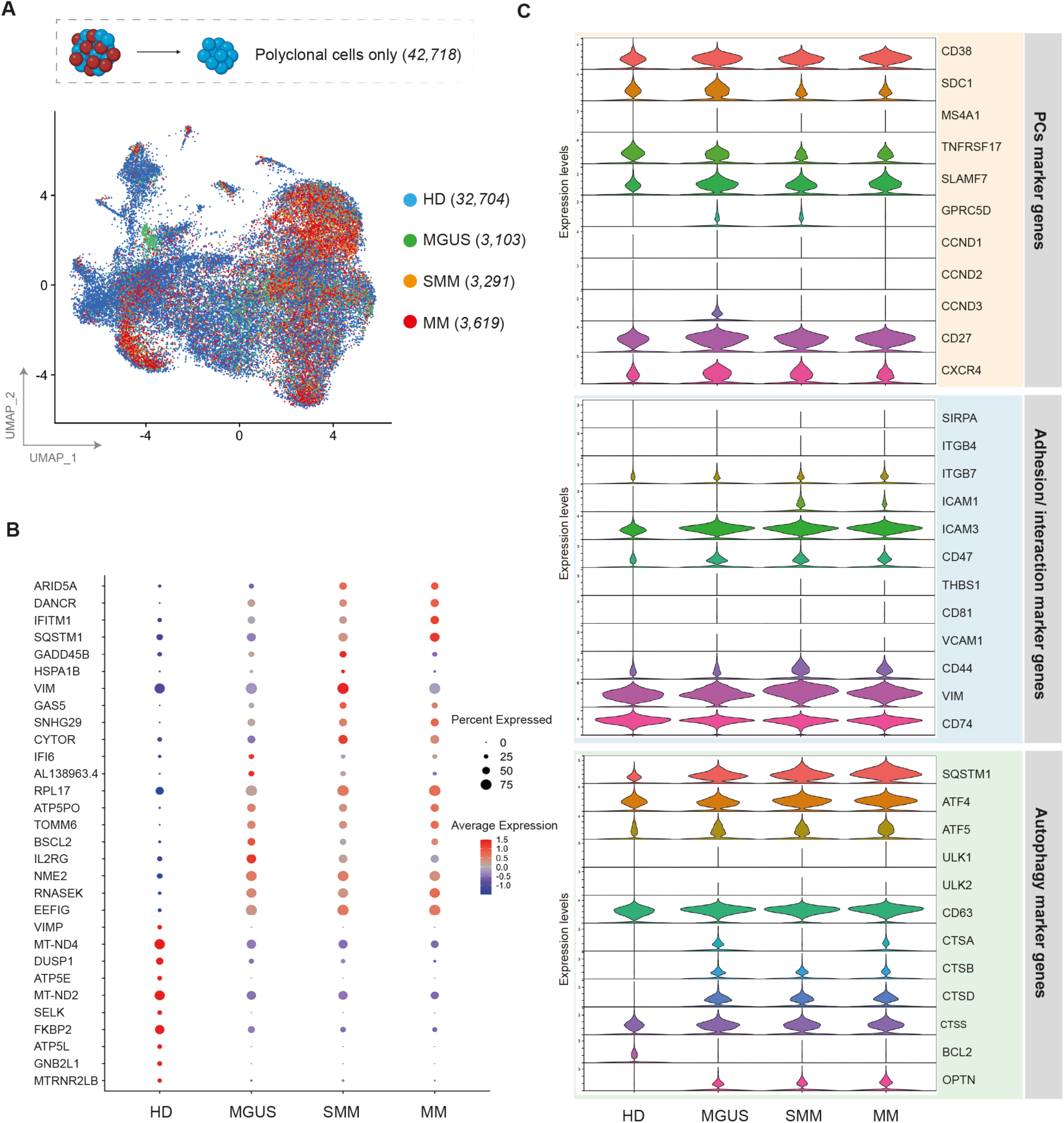
Polyclonal cell selection and marker genes expression by clinical stages. **(A)** UMAP of scRNA-seq data of selected pPCs with colors indicating clinical clusters. Numerical values are reported in the legend for each category. Color legend of polyclonal cells as in Figure 1C. **(B)** Dot plot displaying the top ten marker genes that distinguish each clinical state. The X-axis lists the clinical category, while the Y lists gene names. Circle size corresponds to the number of cells in the category expressing the gene of interest, while shade correlates with the level of expression. **(C)** Violin plot showing expression pattern of selected PCs, adhesion/ interaction and autophagy marker genes across all the clinical stages.

**Figure 3.**
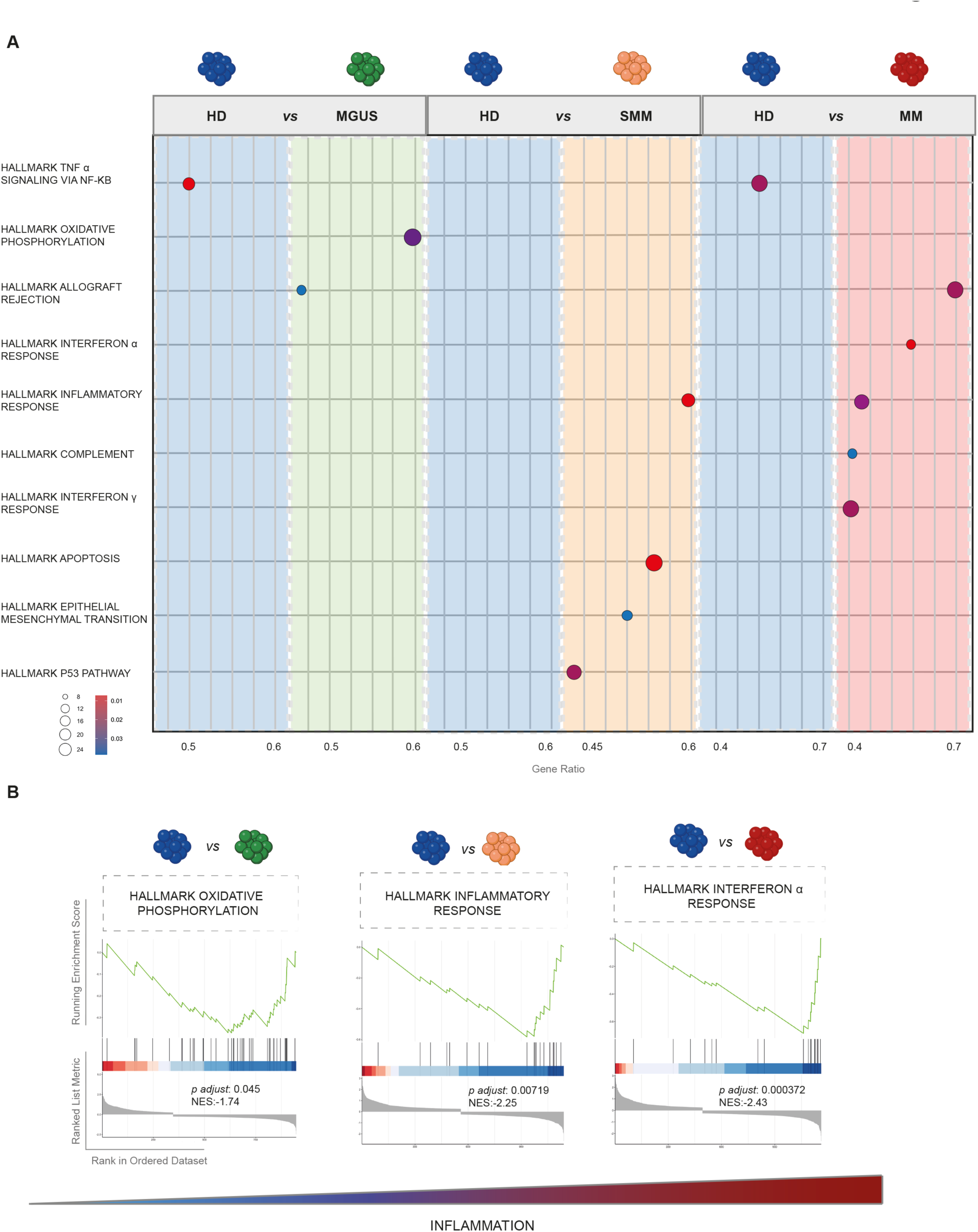
Transcriptional characterization of pPCs. **(A)** Cartoon depicting the deregulated expression profiles of pPCs, derived from 1:1 DE analyses by Wilcoxon test. Circle size corresponds to the number of cells in the category expressing the gene of interest, while shade correlates with the level of expression. Color code as in Figure 1C. **(B)** Gene set enrichment analysis (GSEA) on the genes ranked by their contribution to hallmark oxidative phosphorylation, inflammatory and interferon alpha response.

### Single-cell transcriptomic correlates of immunoparesis

Production of polyclonal immunoglobulins in patients with plasma cell dyscrasias is impaired to variable extents. Particularly, reduction in the production of uninvolved immunoglobulins is seen as a risk factor for progression in asymptomatic conditions^31,40^, and as a risk factor for survival and infections in MM, where it also predicts lower response to vaccination^41,42^. Our unique dataset allowed us to specifically characterize the transcriptional profiles of pPCs in samples with and without immunoparesis, defined as the decrease of one or more of the uninvolved immunoglobulin classes^31^. Having shown the presence of strongly enriched inflammatory pathways in pathological pPCs, we next sought to define their expression profile in immunoparesis-positive patients.

Of n= 46 samples (n=*10,013* pPCs) (**Figure 4A)**, 38 showed immunoparesis. We first observed that pPCs frequency tended to decline in patients showing immunoparesis (**Figure S9A-D**), likely explaining at least in part the decreased production of immunoglobulins. We next analyzed their gene expression profile. The immunoparesis-negative patients showed the expression of genes related to NF-KB (*i.e*. *BIRC3*), cell adhesion (*i.e. CXCR4, SDC1* and *PECAM1*), autophagy (*i.e. FOS, CST3*) and anti-apoptotic (*i.e. DYNLL*1) pathways. As compared to pPCs in patients without immunoparesis, pPCs in patients with immunoparesis showed upregulation of genes related to the interferon pathway (*i.e*. *IRF1, MX1* and *ISG15*), Unfolded Protein Response (UPR) (*ERN1*), cell proliferation and apoptosis (*i.e. H3F3B* and *NOP53*), implying a more deregulated transcriptome (**Figure 4B)**. Then, to explore the functional consequences of gene expression deregulation we performed DE analysis comparing the immunoparesis status within the subset of symptomatic patients (n=23). Here, in presence of immunoparesis pPCs were functionally enriched for interferon related pathways (hallmark IFN α and hallmark IFNγ) and oxidative phosphorylation **(Figure 4C-D).** Taken together, our results reveal consistent changes in expression profiles of pPCs in immunoparesis-positive patients and correlate the presence of a pro-inflammatory phenotype with decreased immunoglobulin production.

**Figure 4.**
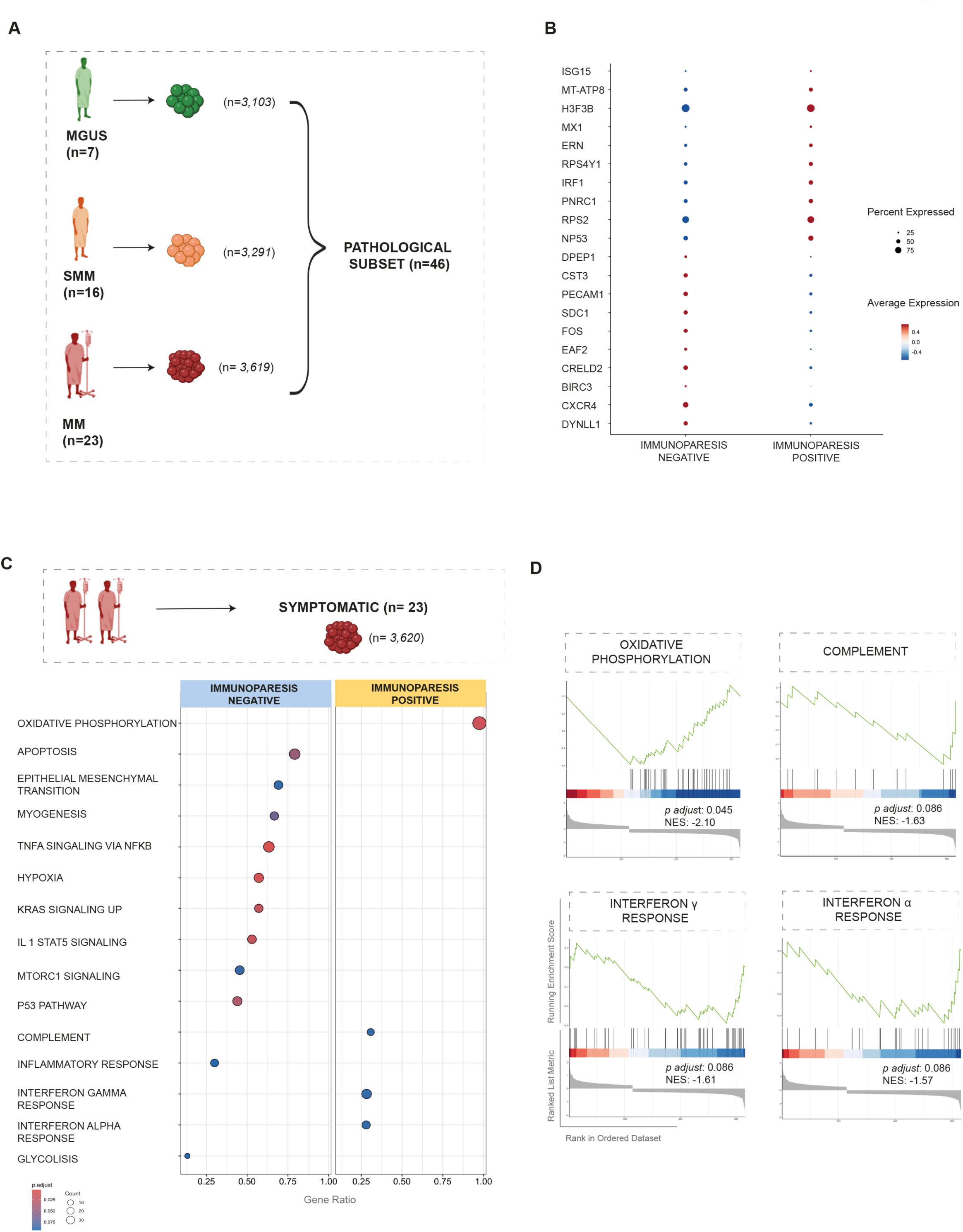
Transcriptomic landscape of immunoparesis. **(A)** Cartoon showing the pathological subset of patients (MGUS, SMM and MM), for downstream DE analyses. Color code as in Figure 1C. **(B)** Dot plot of top ten marker genes that distinguish immunoparesis-negative and immunoparesis-positive patients. The X-axis lists the clinical category, while the Y lists gene names. Circle size corresponds to the number of cells in the category expressing the gene of interest, while shade correlates with the level of expression. Light blue: immunoparesis-negative patients, yellow: immunoparesis-positive patients. **(C)** DE analysis comparing the immunoparesis-negative and immunoparesis-positive patients within symptomatic subset of patients, using Wilcoxon rank sum test, with *padjust* pBenjamini-Hochberg correction. Color code as in Figure 4B. **(D)** Gene set enrichment analysis (GSEA) on the genes ranked by their contribution to hallmark oxidative phosphorylation, complement, interferon alpha and interferon gamma response

### The interactome landscape of immunoparesis in polyclonal PCs

Since progression and persistence of MM malignancy are influenced by the BM TME, and functional properties of both cPCs and pPCs are clearly dependent on interactions with the TME, we next investigated the cell-cell interactions between PCs repertoire and TME in the BM. To gain a first insight into the composition of the TME, we performed scRNA-seq of n=31 paired CD138 positive-CD138 negative cell fractions within our cohort. In total, we integrated *n= 171,928* cells into a single dataset and used the MultiNicheNet pipeline^43^, a tool that infers cell-cell interactions based on ligand-receptor expression and on the pathway activation downstream of the ligand in different cell types. To untangle the complexity of the TME, cPC n= *102,194*, pPC n= *7,447*, and TME cells n=*62,287* were analyzed (**Table S3**). The TME and the PCs’ compartment mapped separately in UMAP plots (**Figure 5A**). Here, we found, n= *148,006* cells of n=23 (*n=89,037* cPCs, n=*3923* pPCs, n=*55,046* TME) characterized by immunoparesis and n= *23,922* cells from n=8 immunoparesis-negative samples (n= *13,157* cPCs, n= *3,524* pPCs, n= *7,241 TME*) (**Figure 5B**). Splitting the TME data by cell type, n=*32,966* T cells, n=*3,472* B cells, n=*15,261* myeloid cells, n= *6,589* NK cells, n=*1,019* dendritic cells and n=*2,980* HSCs were analyzed and no significant differences were observed in the relative composition of the TME populations (**Table S3, Figure S10)** between immunoparesis-positive and immunoparesis-negative patients **(Figure 5B, Table S3, Figure S10)**. As expected, we found a significant depletion of pPCs in immunoparesis-positive patients (*padjust = 0.004*)(**Figure S10**).

**Figure 5.**
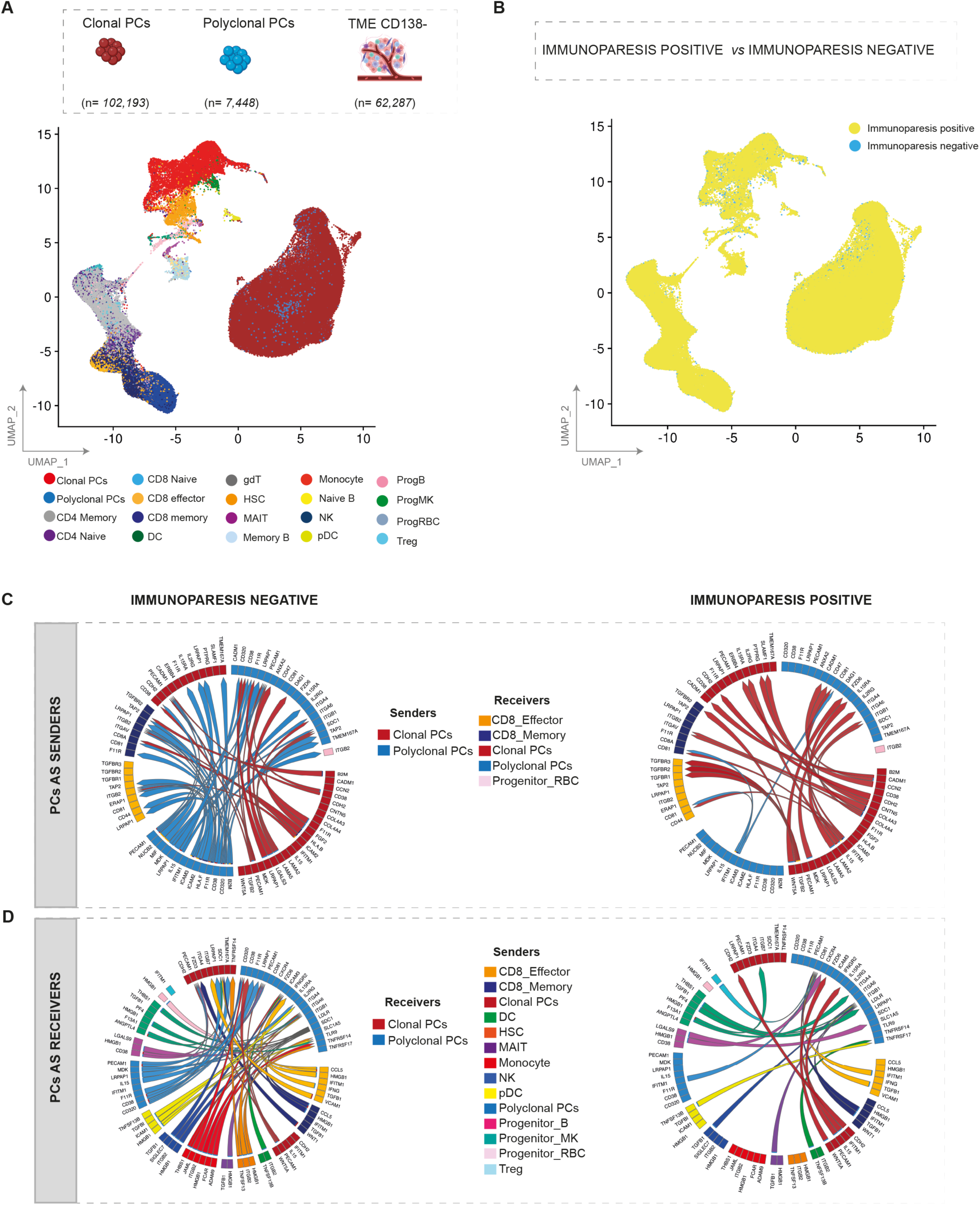
Predicted interactions of PCs with microenvironment. **(A)** UMAP representation of PCs integrated with TME, colors indicate TME cell types. **(B)** UMAP plot of PCs integrated with TME colored according to immunoparesis. Color code as Figure 4C. **(C)** Circos plots illustrate ligand-receptor Interactions between pPCs (blue), cPCs (red) and TME, according to immunoparesis clinical status. Ribbon arrows indicate directionality of communication, from sender to receiver populations, while the arrow’s color signifies the specific sender cell type expressing the ligand. Upper panel: PCs were set as sender. Lower panel: PCs were set as receiver.

To understand disruption of inter-cellular relationships in this scenario, we first analyzed the immunoparesis-negative setting. Comparing top-ranked ligand-receptor interactions and analyzing the downstream targets, pPCs were predicted to be able to send signals to specific cell compartments such as pPCs, CD8-effector, CD8 memory and progenitor RBC cells **(Figure 5C, Figure S11A-B)**. Mining PCs as senders, of relevance were the *CD38: PECAM1* (CD31) cell interaction –with a role in regulation of cell migration-, and a broad spectrum of interactions mediating cell adhesion involving *ICAM2, ICAM3* and *ITGB2* (T lymphocytes), already described in promoting cell survival processes.^44–48^ Furthermore, cPCs and pPCs, displayed the ability to receive signals from myeloid (DCs, HSCs, monocytes, pDC and Mk progenitor cells, lymphocytes (CD8 effectory, CD8 memory, NK, progenitor B and MAIT cells), compartments. In this scenario, immune and inflammatory responses were driven by the specific interaction of THBS-1 (in monocytes and Mk progenitor cells) with *SDC1* (in pPCS) and *ITGA6* (in pPCS), as well as by the signaling of *HMGB-1* (in HSCs, NK, pDCs, MAIT, CD8 cells, progenitor B cells and monocytes): *SDC1* (in pPCS). In addition, we found the *CCL5:SDC1* ligand-receptor interactions between T cells and cPCs/pPCs, known to be involved in tumor cell migration^49^. Looking at the downstream deregulated gene targets, derived by ligand-target correlation, genes implicated in the autophagy axis and in ER stress (*BNIP3, MT1X, MT2A*)^50,51^ were upregulated as well as B cell survival pathways regulators (*i.e. CD74* and *ITGA6*)^52^ (**Figure S12A-B**). These findings highlight an active involvement of pPCs in the pro/inflammatory TME in immunoparesis-negative cell landscape. However, pPCs in this setting maintain their functional role through autophagy, ER stress regulation and B cell survival pathway activation.

Conversely, cPCs became the main contributor of TME interactions in the immunoparesis-positive cell landscape, where a significant depletion of pPCs:TME interactions was observed (**Figure 5D, Figure S13A,B)**. These differences were not simply determined by a depletion of pPCs since the differential expression analyses are normalized for cell count.

In detail, cPCs and pPCs were characterized by the inflammatory signaling mediated by the *IFITM1:CD81* interaction (pPCs: pPCs, pPCs:CD8 effector/memory cells, Treg:pPCs), that is involved in IFN-gamma response pathway^53,54^. Of note, in this scenario where the interactions between myeloid compartment and pPCs were significantly depleted, DCs and pDCs were found to sustain pPCs survival through *BAFF-BCMA* axis^55–57^.

Furthermore, the axis *TGFB2:TGFBR1/TGFBR2/ TGFBR3* mediated interactions between cPCs and CD8 T cells together with *TGFB1:ITGB1* (MAIT and NK with pPCs), suggesting a role of PCs in the immunosuppression of T-cell compartment. Of note, the γ-chain cytokine *IL-15* was shown to signal from pPCs in an autocrine manner to pPCs and in a paracrine manner to cPCs by binding the common cytokine receptor γ chain (γc or *CD132*; encoded by the *IL2RG* gene) in the immunoparesis-negative setting. Conversely, in immunoparesis-positive patients, this cytokine signals from cPCs to cPCs and pPCs through the *IL15RA* receptor^58–61^. Only in this latter setting, the downstream signalling involves activation of pro-inflammatory pathways. Furthermore, downstream target genes related to inflammation and interferon pathways were upregulated (i.e. *IFI16, IFIT3, PLSCR1*)^62^ as well as negative regulators of NF-kB pathway such as *TNFSF10*. Conversely, in this setting genes related to autophagy and B cell maintenance resulted downregulated (i.e. *CSTH, BNIP3, MTX1, MT2A*)^34,50^ (**Figure S14A,B)**. Altogether these data support the presence of a pro-inflammatory status in immunoparesis-positive patients, driven by specific cellular interactions that are disrupted in immunoparetic patients likely impacting pPCs survival and function.

### Signature analysis allows prediction of survival in independent datasets

The presence of residual PCs has been correlated with prognosis in MM patients^33,34^. Our data show that even pPCs in patients can be profoundly dysregulated and different from PCs in HDs. Our unique dataset therefore allowed us to generate a “healthy PC” (hPC) signature through the union of the differentially expressed genes in the three contrasts of PCs from HD with pPCs from MGUS, SMM, and MM patients respectively. Importantly, our hPC signature corrects for the dysregulated expression of pPCs in patients and incorporates a total of *n=148* genes (**Figure 6A**, **Table S4**) representing genes truly expressed in hPCs only. Then, a specific immunoparesis signature, composed by n=10 deregulated genes, was derived by comparing immunoparesis-negative with immunoparesis-positive patients, in asymptomatic and symptomatic subset (**Figure 6B**, **Table S5**).

**Figure 6.**
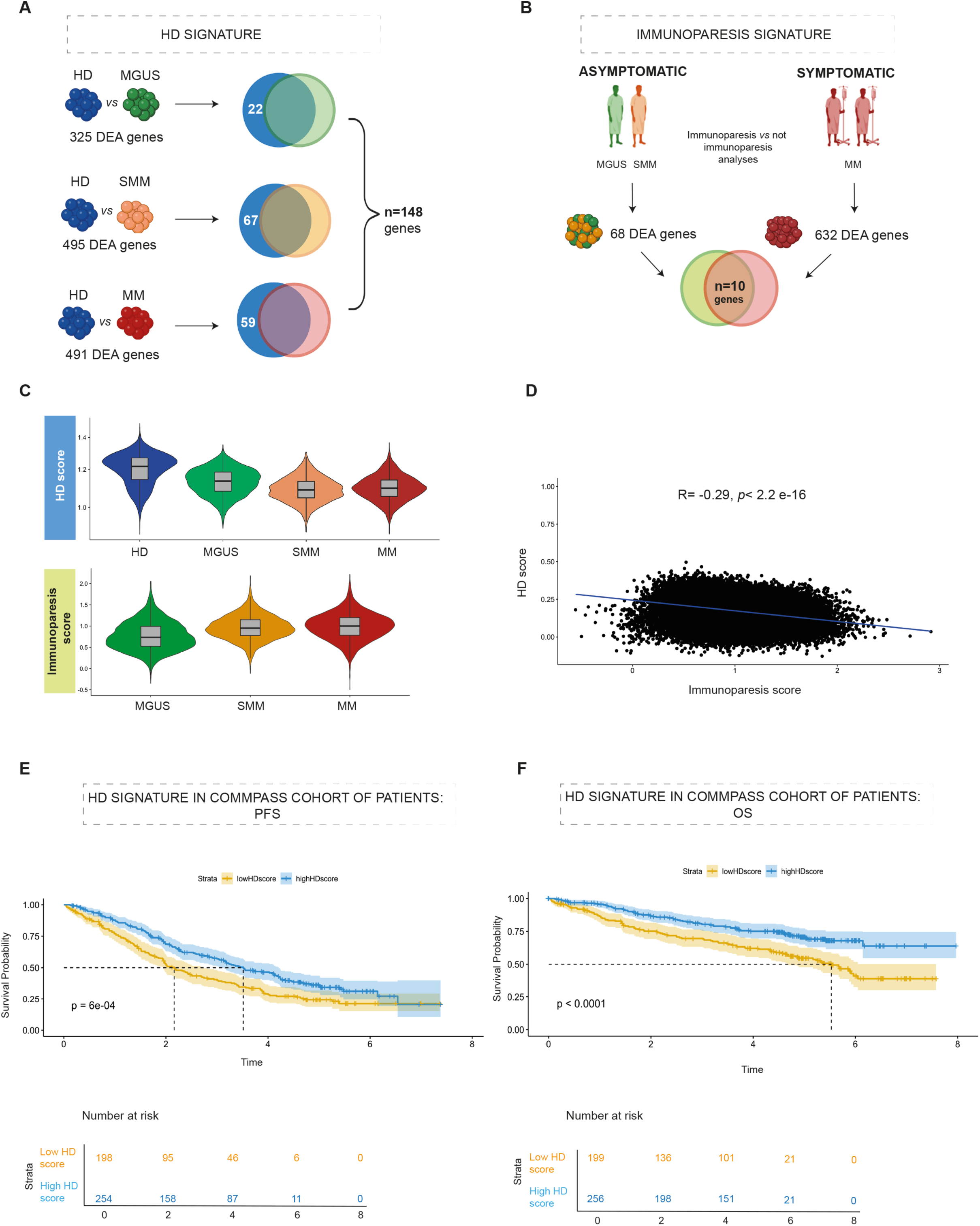
A «hPC» signature analysis allows prediction in independent datasets. **(A)** HD gene signature defined by a DE analysis between HDs and MGUS, SMM and MM. For each contrast genes significantly upregualted in HDs (Benjamini—Hochberg adjusted p.value < 0.05, logFoldchange > 0.1) were retained. Venn diagram to show the intersection of deregualted genes within each comparison. Color code as Figure 1D. **(B)** Immunoparesis gene signature defined by a DE analysis between immunoparesis-negative and immunoparesis-positive patients. Venn diagram to show the intersection of deregualted genes. Color code as Figure 4B. **(C)** Violin plot showing the inference of HD and immunoparesis signatures in the cohort of patients. Color code as Figure 1D. **(D)** Scatterplot of HD and Immunoparesis signatures: R: – 0.29, *p.value* < 2.2e^-16^. **(E, F)** Clinical impact of HD signature in CoMMpass RNAseq data. Progression-free survival (PFS) **(E)** and overall survival (OS) **(F)** Kaplan-Meier curves in the cohort of patients.

Applying the hPC signature in our cohort, a gradient could be observed whereby the signature was enriched in HD, and progressively less represented in pPCs from MGUS, SMM and MMs cases (**Figure 6C, upper panel**). The same inference was performed with the immunoparesis signature which showed an opposite trend, resulting more enriched from MGUS to MMs (**Figure 6C, lower panel**). Indeed, the hPCs signature anti-correlated with the signature of immunoparesis in our pPC dataset, implying that cases enriched for abnormal pPCs have a lower propensity to produce polyclonal immunoglobulins and might express a more corrupted phenotype than healthy PCs (R: – 0.29, *p.value* < 2.2e^-16^, **Figure 6D**).

Finally, we decided to determine if enrichment for the hPC signature could have a clinical impact within a NDMM population. We hypothesized that in bulk RNAseq generated from CD138-selected cells part of the signal could be generated by pPCs. We therefore interrogated the RNAseq data of CoMMpass and ranked patients based on the fractional representation of the hPCs signature. A simple categorical distinction based on the enrichment of the hPCs signature above and below the median clearly separated patients based on their survival. Specifically, the median progression-free survival (PFS) was 3.52 years [95% confidence interval, 3.13-4.21 years] in patients above the median and 2.16 years [95% confidence interval, 1.94-2.75 years] in patients below the median (*p.value* = 0.0006, log-rank test, **Figure 6E**). Median overall survival (OS) was not reached in patients above the median and was 5.5 years [4.68-not reached years] in patients below the median (*p.value* < 0.0001, log-rank test, **Figure 6F**). Based on these results and assuming that the hPCs score might represent a surrogate of residual functional “polyclonality” within clonal PCs, we could argue that the maintenance of a normal polyclonal population impact on patient’s survival, potentially explaining the observed correlation between immunoparesis and prognosis^42^.

## DISCUSSION

The TME architecture has long been implicated in initiation and evolution of PC dyscrasias. pPCs are an obvious component of the TME but have been understudied despite a clear biological and clinical relevance. Part of this depends on technical issues. For example, pPCs cannot be separated from cPCs based on isolation with magnetic beads nor with flow-sorting. In this scenario, we applied a comprehensive approach including sc-RNA sequencing coupled with BCR genotyping to unambiguously identify pPCs in patients with plasma cell dyscrasias. To confirm the robustness of our approach, CNA analysis on pPCs and cPCs showed clonal aneuploidies only in the latter population.

Using this advanced approach, we provide evidence that pPCs would not necessarily be identified with accuracy based on transcriptomics alone. Indeed, even after removing immunoglobulin genes, i.e. highly expressed cell-specific genes accounting for a big part of the transcriptomic diversity between cells, RNAseq alone may not effectively separate clonal and polyclonal cells^33,34^. Our data showing that pPCs did not necessarily segregate away from cPCs from the same patient in the UMAP space, suggest they are themselves phenotypically aberrant. This is different from previous studies^34^, and can be explained at least in part by the innovative methodology we employed.

pPCs from patients showed the expected differential expression of canonical marker genes as compared to cPCs. However, we also convincingly show that GPRC5D expression is natively lower in pPCs than in cPCs, a finding which may suggest that anti-GPRC5D bi-specific antibodies may relatively spare pPCs and explain the lower prevalence of infections in such patients contrary to anti-BCMA antibodies^63^.

However, aside from canonical marker gene expression, our data reveal a profoundly dysregulated transcriptomic landscape of pPCs from patients affected by asymptomatic or symptomatic PC dyscrasias when compared to BM PCs derived from HDs sourced with a broad age range. In this setting, pPCs showed upregulation of *IFITM1, SQTM1*, *CTSB*, *CTSD*, *OPTN*^53,54^ markers as well as *ICAM1*^64–68^, implying an upregulation of inflammation, autophagy and interferon pathways, with a climax from asymptomatic to symptomatic patients. Of note, the primary purpose of autophagy is to sustain the cellular homeostasis in extreme conditions^69^, such as the ones promoted by oxidative and proteotoxic stress due to an intense production and secretion of immunoglobulins^69–71^. Therefore, autophagy is fundamental for long-lived PCs to optimize the protein metabolism and the cell viability^70–73^ and we believe that this pathway is exaggerated in the setting of residual pPCs with the BM of patients with PC dyscrasias. Our findings also have translational implications. pPCs derived from immunoparesis-positive patients showed upregulation of *IRF1, MX1* and *ISG15, ERN1, H3F3B* and *NOP53* genes suggesting activation of IFNα and IFNγ pathways, UPR and cell proliferation/apoptosis signaling in the suppression of uninvolved immunoglobulins. We further extended this analysis by looking at intercellular interactions between PCs and the TME within each patient. Here, we provide evidence of a TME poised for inflammation in immunoparesis-positive patients. In this scenario, we observed a significant decrease in the number of interactions of pPCs with other immune cells in the TME. Furthermore, pPCs were characterized by a pro-inflammatory transcriptome. The interaction between IFITM1 and CD81 and the activation of the TGFB2:TGFBR1/TGFBR2/TGFBR3 axis that we see in our patients were indeed shown to be mainly driven by IFN pathways and may affect pPCs survival and function, including production of IgM and IgG^74^.Furthermore, at the cell-intrinsic level, our data showed a reduction in expression of authopagy marker genes combined with an increase of pro-apoptotic genes. Altogether, our findings shed new light on mechanisms of deregulated pPCs homeostasis leading to immunoparesis in patients affected by PC dyscrasias. Furthermore, based on previous studies^75–77^ we speculate that in these conditions the immune microenvironment could be less competent in antigen-presenting activity as well as in immunosurveillance, further adding to the complex interrelationships between antibody production, immunity and tumor progression.

One important question is what the primary cause of an inflamed TME is. Disease pathogenesis and our data may suggest that cPCs, through perturbed interaction with pPCs and other cells in the TME, might be responsible. However, it can also be argued that a perturbed and a pro-inflammatory TME can itself has a primary driver role in PC dyscrasia development, deranging the normal BM PC transcriptional program and thus promoting the growth of pre-existing cPCs –harboring initiating genomic lesions-whose proliferative potential was not expressed until then. The notion that cPCs can progress to an overt MM without acquiring new genomic events and thus potentially based on microenvironmental stimuli, is indeed quite substantial^20,73^ and our data shed some light for the first time on the type of selection applied from the TME on BM polyclonal PCs.

Intriguingly, we also identified a specific “healthy” PC signature by comparing PCs from HD with pPCs of patients rather than to total CD138+ cells, that resulted decreased across disease-stages and anti-correlated with the immunoparesis signature. Notably, using RNAseq data from the CoMMpass dataset, patients with an HD signature enrichment over the median showed a better PFS and OS. These findings emphasize the role of residual functional “polyclonality” within BM PCs, suggesting that the maintenance of a normal polyclonal population impacts on patient’s survival and potentially explaining the observed inverse correlation between immunoparesis and prognosis ^75^. In conclusion, our study employs a new methodology of analysis to gain deep insights into transcriptional landscape of pPCs in PCs dyscrasias, suggesting they are key players in the TME, and their disruption has biological and clinical consequences whose understanding may impact research and treatment of MM.

## ACKNOWLEDGMENTS

This work was supported by the European Research Council under the European Union’s Horizon 2020 research and innovation program (grant agreement No. 817997), by International Myeloma Society (IMS) and Paula and Rodger Riney Foundation Translational Research Grant Application-2023 and by Associazione Italiana Ricerca sul Cancro (IG25739). AIRC IG 24365 to AN. M.C.D.V. was funded by Umberto Veronesi Foundation and by Pfizer Global Medical Grants (grant tracking No. 75340503). This study was supported in part by Italian Ministry of Health—Current research IRCCS.

## AUTHOR CONTRIBUTIONS

**N.B.**, **M.C.D.V. F.L.,** conceived the project; **N.B., M.C.D.V., F.L.,** designed the experiments; **N.B., M.C.D.V., C.D.M, L.P.,** enrolled and followed up patients and acquired data; **M.C.D.V., F.L., A.M., A.M.,** performed single cell experiments; **A.M.** verified the raw data before analyses and performed pre-processing bioinformatic analysis; **M.C.D.V., F.L., A.M.,** performed the bioinformatic analysis and interpreted data; **M.L., S.P., S.F., S.L., M.S., N. L., G. C., A.N., F.T., K.F., T.C., F.P.,** contributed with scientific discussions; **A.C., M.T.,** performed the flow cytometry experiments; **N.B., F.L., A.M., M.C.D.V.,** wrote the manuscript, which has been revised by all the authors.

### Declaration of interests

N.B. received honoraria for Amgen, GSK, Janssen, Jazz, Pfizer, Takeda.

M.C.D.V. served as Advisory Board for Takeda, and on Speakers Bureau for Janssen, Sanofi and GSK.

F.P. received honoraria during the last two years for lectures from Novartis, Bristol-Myers Squibb, Abbvie, GSK, Janssen, AOP Orphan and for advisory boards from Novartis, Bristol-Myers Squibb/ Celgene, GSK, Abbvie, AOP Orphan, Janssen, Karyiopharma, Kyowa Kirin and MEI, Sumitomo, Kartos. The remaining Authors declare no competing financial interest.

## STAR * METHODS

KEY RESOURCES TABLE.

| REAGENTS or RESOURCE | SOURCE | IDENTIFIER |
| --- | --- | --- |
| **Critical commercial assays** |  |  |
| BCR Amplification Kit | 10X Genomics | PN-1000253 |
| Chromium Next GEM Single Cell 5' Reagent Kits v2 | 10X Genomics | PN-1000263 |
| Chromium Next GEM Chip K | 10X Genomics | PN-2000286 |
| Sample Index PCR | 10X Genomics | PN-1000190 |
| Dual Index Kit TT Set A | 10X Genomics | PN-1000215 |
| SPRIselect beads | SPRIselect Reagent Kit | Cat# B23318 |
| High Sensitivity D5000 ScreenTape | Agilent | Cat# 5067-5592 |
| High Sensitivity D5000 Reagents | Agilent | Cat# 5067-5593 |
| **Software and algorithms** |  |  |
| ggplot2 | CRAN | v 3.4.4 |
| R | CRAN | v 4.3.2 (2023-10-31) |
| Seurat | CRAN | v 4.4.0 |
| InferCNV | Bioconductor | v 1.16.0 |
| MultiNicheNet | GitHub repository | V 1.0.3 |
| ClusterProfiler | Bioconductor | v 4.10.0 |
| DoubletFinder | GitHub repository | v 2.0.4 |
| Harmony | CRAN | v.1.2.0 |
| Survival | CRAN | v 3.5-8 |
| Survminer | CRAN | v 0.4.9 |
| ComplexHeatmap | Bioconductor | v 2.18.0 |
| **Antibodies and reagents** |  |  |
| EasySep™ CD138 Positive selection kit II | STEMCELL Technologies | Cat# 17887A |
| BamBanker | NIPPON Genetics Europe | Cat# BB02 |
| RPMI Medium 1640 + GlutaMAX™ | gibco | Cat# 61870-010 |
| Fetal Bovine Serum, FBS | gibco | Cat# A5256701 |
| Phosphate buffered saline, PBS | gibco | Cat# 70011-044 |
| UltraPure™ 0.5M EDTA, pH 8 | Invitrogen | Cat# 15575-038 |
| Ficoll-Paque™ PLUS | Cytiva | Cat# 17144002 |
| Erythrocyte lysing solution | Bio-Techne (Cytognos) | Cat# CYT-BL |
| Quant-iT™ PicoGreen™ dsDNA Assay kit | Invitrogen | Cat# P7589 |
| **Visualization softwares** |  |  |
| Illustrator 2024 | Adobe |  |
| Photoshop 2024 | Adobe |  |
| Prism 10 | Graphpad | v 10.1.1 |
| Biorender |  |  |
| **Databases** |  |  |
| GEO accession number<br>#GSE193531 |  |  |
| GEO accession number<br>#GSE230705 |  |  |

## RESOURCE AVAILABILITY

### Lead contact

Further information and requests should be directed and will be fulfilled by the lead contact, Niccolò Bolli.

### Materials availability

This study did not generate new unique reagents.

### Data and code availability

The raw data and processed scRNA-seq data, including gene expression matrices and cell annotations, have been deposited at the European Genome-Phenome Archive (https://ega-archive.org/) under accession numbers EGAC50000000232.

R script written for the analysis of the single cell datasets is available on https://gitlab.com/bollilab. ScRNA-seq data and R script required to reanalyze the data reported in this manuscript, are available from the lead contact request.

## EXPERIMENTAL MODEL AND STUDY PARTECIPANTS DETAILS

### Sample selection

For this study, bone marrow (BM) aspirates from of n= 46 samples, affected by MGUS (n=7), SMM (n=16) and MM (n=23) were obtained at Fondazione IRCCS Ca’ Granda Ospedale Maggiore Policlinico, Milan (Italy), according to IMWG criteria (PMID: 25439696). Detailed clinical information for each case is provided in **Table S1.** The study was conducted in accordance with the Declaration of Helsinki and approved by the Institutional Review Board (Ethical Committee # 553_2022). Written informed consent was obtained from the patients, as appropriate. All patients were de-identified prior to processing.

One sample from healthy donor (HD) was collected at Fondazione IRCCS Ca’ Granda Ospedale Maggiore Policlinico, Milan (Italy), as well as (n = 3) from Erasmus MC Cancer Institute, Rotterdam, (The Netherlands). N=14 (HDs) samples were obtained from publicly available databases^34,36^. In particular, from Boyarsky et al^34^, a subset with only count derived from the 9 healthy donors was performed (NCBI Gene Expression Omnibus (GEO) database accession number #GSE193531). From Duan^36^ et al., data derived from 5 healthy donors have been used as control: within this, only counts derived from sorted cell population B (CD19+/CD138+/CD38+) and cell population D (CD19-/CD38+/CD138+) were used for further analysis (GEO accession number #GSE230705). The range of HD ages is 25-79 years. The polyclonal status of the healthy donor collected in our institution was defined through scV(D)J seq defining the absence of PCs clones with a frequency higher than 5% of the entire PCs population.

### Sample processing

Mononuclear cells (MNCs) were isolated by Ficoll-Paque medium (Cytiva, Cat# 17144002) density gradient centrifugation. CD138+ BM cell fractions were isolated by immune-magnetic approach through positive selection with the EasySep™ CD138 Positive selection kit II (STEMCELL Technologies, Cat#17887A), after erythrocyte lysis (Cytognos, Cat# CYT-BL), following manufacturer’s instructions. Cells were counted based on Trypan Blue staining and the resulting cell suspension was used for downstream applications.

N=8 CD138-samples were cryopreserved in liquid nitrogen using Bambanker (NIPPON Genetics Europe, Cat. #BB02).

### Thawing of Cells

Cell thawing was conducted following standard protocols. N=8 samples, stored in liquid nitrogen, were swiftly transferred to a 37°C water bath until the vial contents were fully dissolved. Subsequently, the content was transferred to a Falcon tube containing pre-warmed RPMI (Cat# 61870-010) containing 10%FBS (Cat# A5256701), followed by a washing step.

## Method details

### Single-cell RNA 5’ sequencing library construction and sequencing

Isolated cells from each sample were washed three times with 1X Phosphate-Buffered Saline (PBS, calcium and magnesium free) containing 0.04% weight/volume FBS (Gibco, Cat# 70011-044) and manually counted by the Burker chamber. In order to have a desired recovery target of 9,000 cells, the volume of cells to load was calculated for each sample according to “Cell Suspension Volume Calculator Table” on Chromium Next GEM Single Cell 5’ v2 protocol (CG000330 Rev E). Then Single-cell 5ʹ RNA-seq libraries were generated using the Chromium Next GEM Chip K (10X Genomics; PN-2000286) with Master Mix from Chromium Next GEM Single Cell 5’ Reagent Kits v2 (Dual Index) (10X Genomics; PN-1000263), Single Cell VDJ 5’ Gel Beads (10X Genomics; PN-1000253) following standard manufacturer’s protocols. Briefly, cell suspensions were loaded onto the Single Cell K chip followed by gel beads in emulsion (GEM) generation on the Chromium Controller. Next, GEMs were broken, and pooled fractions were recovered.

Following the barcoded, full-length cDNA amplification, the cDNA was purified by SPRIselect Reagent (Cat# B23318) and quantified using Picogreen™ dsDNA assay kit (Cat# P7589) and quality assessed using High Sensitivity D5000 ScreenTape (Agilent, Cat# 5067-5592) with High Sensitivity D5000 Reagents (Agilent, Cat# 5067-5593) at the Agilent 4150 TapeStation machine. Then, Illumina R2 sequence, as well as P5, P7, i5 and i7 sample indexes were added according to standard protocol procedure (10X Genomics; Library Construction Kit, PN-1000190 and Dual Index Kit TT Set A, PN-1000215, respectively). Parallelly, BCR Amplification Kit (10X Genomics; PN-2000253) was used for V(D)J analysis. High Sensitivity D5000 ScreenTape and its reagents were used to assess the library quality, as well as Picogreen™ to determine libraries concentration. Finally, Generated libraries were combined according to Illumina specifications and sequenced on a NovaSeq6000 platform (Illumina sequencing system) using a 150bp Paired End protocol, targeting approximately 100,000 reads/cell and 15,000 reads/cell for 5’ gene expression and V(D)J enriched libraries, respectively, according to manufacturer’s recommendations.

### CD138+ pre-processing single-cell RNA sequencing analysis

The gene expression (GEX) and V(D)J single-cell RNA-seq data were aligned and quantified using the Cell Ranger (v 7.1) pipeline (https://www.10xgenomics.com/) against the human genome GRCh38 (refdata-gex-GRCh38-2020-A and refdata-cellranger-vdj-GRCh38-alts-ensembl-5.0.0, respectively) using the functions “counts” and “vdj counts” for single-cell RNAseq and V(D)J deconvolution, respectively. Then, Seurat pipeline (v 4.4.0,) was used to analyze expression matrices. Low-quality cells defined as mitochondria gene content >5%, or number of detected genes < 200 or > 3,000 were removed using Seurat pipeline (v 4.4.0, https://satijalab.org/seurat). In order to avoid cell clustering confounding factors, immunoglobulin related genes, were removed for downstream analyses. Doublets were filtered out using DoubletFinder^78^ (v 2.0.4) with the core statistical parameters (nExp, pN and pK) determined automatically using recommended settings for each sample. nCounts_RNA, n_FeatureRNA and percent_MT were checked for each sample and reported in **Figure S6 A, B**. An automated cell assignment was applied on previously filtered single-cell which were projected onto a complete Bone Marrow reference map (v4.4.0, https://satijalab.org/seurat/articles/multimodal_reference_mapping.html)^79^. The clonotype information was incorporated for each cell by matching the barcodes from the single-cell V(D)J (scVDJ) with those from the single-cell gene expression (scGEX). This process facilitated the classification of cells as either harboring clonal or polyclonal clonotypes.

### Polyclonal cell selection

In order to analyze a clear population of genotypically identified polyclonal plasmacells (PCs), a strict multistep process was applied: *(i)* for each patient the dominant clonotype was defined as the recurrent V(D)J rearrangement within the clonal population; *(ii)* the nucleotide sequence of the CDR3 of any non-clonal clonotype (heavy chain and light chain rearrangement) was blasted against the nucleotide sequence of the dominant V(D)J rearrangement; *(iii)* all those cells whose the blast analysis return with certain grade of homology were manually inspected. Cells with a dubious homology (more than 90% as for the IGH as for IGK/IGL chain) with the dominant clonotype were removed, as well as cells sharing only the light chain rearrangement with the dominant clone were considered “light chain escape derived clones” and categorized within the clonal PCs. *(iv)* Samples with an higher degree of homology between cell’s rearrangements underwent to a phylogenetic tree analysis through the Immcantation pipeline^80,81^ to exclude any phylogenetic correlation between the dominant clonotype and other putative polyclonal PCs; *(v)* Each sample underwent to inferCNV analysis (https://github.com/broadinstitute/inferCNV, (see detailed methods below)^37^ to dissect at transcriptional level the different copy number variants (CNV) background between the clonal and polyclonal PCs populations. Each sample analysis was performed using the same healthy donor PCs as normal control (a complete overview of CNVs is reported **Figure S1-S4**); *(vi)* a final list for each sample containing specifically cell barcodes for polyclonal and clonal PCs were created and used for downstream analysis.

### Plasma cells single-cell RNA sequencing analysis

ScRNAseq analysis as performed by Seurat pipeline. Samples were merged in lists based on their clinical status (HDs, MGUS, SMM and NDMM) and counts normalized. Then, the integration was performed using the *FindTransferAnchors* and the *IntegrateData* functions with RPCA method. The integrated Seurat object was scaled, and the most variable features were obtained through the *FindVariableFeatures* function. Based on these genes the principal components analysis was performed through *runPCA* function.

Batch effect correction was applied to PCs using Harmony^82^ (v.1.2.0) with function *RunHarmony* using the origin of the samples as variable for correction. Then, the harmonized matrix was used to plot uniform manifold approximation and projection (UMAP). For the polyclonal BM scRNA-seq data set, 30 principal components (PCs) were used for dimensional reduction and cell clustering. The resolution parameter was 0.1. Differential expression analysis between different clinical stages were performed using FindMarkers function from Seurat package (v 4.4.0). Cluster specific markers were identified by *FindAllMarkers* and *FindMarkers* functions. Violin plots to compare expression across cell types or clinical groups were made using VlnPlot function in R.

Functional enrichment analysis was performed with genes by clusterProfiler package^83^ (v. 4.10.0), for the hallmark gene sets provided by the Molecular Signature Database (MSigDB) (msigdbr, v.7.5.1).

Wilcox test with pvalueCutoff= 0.1 was applied for all analyses. Figures were plotted by ggplot2 (v 3.4.4) in R.

### Single-cell copy number variation analysis

Copy number alterations (CNAs) were inferred from single cell gene expression data using inferCNV R package^37^ (v 1.16.0, https://github.com/broadinstitute/inferCNV), which computes gene expression intensities across genomic positions from malignant cells in comparison to a set of reference cells (for each comparison the same healthy donor have been used for a total of 4153 normal PCs)

The algorithm was run with the following arguments: cutoff=0.1, HMM_type = ‘i3’, cluster_by_groups=TRUE, denoise=TRUE, HMM = TRUE. CNV-level subpopulations were determined using analysis_mode = ‘subclusters’, to partition cells into groups having consistent patterns of CNVs

### Single-cell HD signature definition and scoring

HD gene signature defined by a differential gene expression analysis (“wilcoxauc” R function) between HDs and MGUS, SMM and NDMM. Then, for each contrast only genes significantly upregualted in HDs (Benjamini—Hochberg adjusted p.value < 0.05, logFoldchange > 0.1) were retained. Then, we calculated the intersection of deregualted genes within each contrast though the ComplexHeatmap (v 2.18.0) function *make_comb_mat* using the “distinct”mode to compute only those genes specifically deregulated in each contrast.

Then, the Gene signature activity in single cells was estimated using the *AddModuleScore* function from Seurat (v 4.4.0), and CompleaxHeatmap. Gene sets used for signature scoring are listed in **Table S4.**

To estimate the activity of HD gene expression signature for each sample in the publicly available MMRF bulk RNA-sequencing dataset (IA17 release, https://research.themmrf.org), that includes bulk RNA-seq data from the CD138+ fraction derived from NDMM patients. Survival analysis was performed using survival (v 3.5-8) and survminer (v 0.4.9) packages.

### CD138-pre-processing single-cell RNA sequencing analysis

GEX and V(D)J scRNA-seq data of matched n=31 CD138-samples (n=23 fresh and N=8 frozen) were aligned and quantified using the Cell Ranger pipeline (v7.1) against the human genome GRCh38. For GEX data, the reference used was refdata-gex-GRCh38-2020-A, while for V(D)J data, the reference was refdata-cellranger-vdj-GRCh38-alts-ensembl-5.0.0.

To ensure data quality, cells expressing less than 200 or more than 3000 features, and those expressing over 5% (fresh samples) and 10% (frozen samples) of mitochondrial genes, were filtered out using the Seurat pipeline (v4.4.0).

To identify and remove potential doublets, DoubletFinder^78^ (v2.0.4) was employed with core statistical parameters (nExp, pN, and pK) determined automatically using recommended settings for each sample. Filtered single-cell transcriptomes were then projected onto a comprehensive Bone Marrow reference map using Seurat. Subsequently, cells displaying a clonal VDJ profile, along with all plasma cells, were omitted from the analysis to ensure the derivation of uncontaminated samples representing the tumor microenvironment (TME). All cells derived from the n=31 samples (positive and negative fractions) were integrated following the *FindTransferAnchors* and the *IntegrateData* functions using CCA method.

### Cell-cell communication and interaction analysis

Cell-cell communication analysis was performed using the MultiNicheNet package (version 1.0.3)^43^ implemented in R, to infer potential interactions between cell types using the expression of ligands, receptors, and potential targets within signaling pathways. The MultiNicheNet analysis includes 31 matched samples, it was performed integrating CD138+ cells (comprising both polyclonal and clonal PCs) and CD138-cells for the TME totaling N=171928 cells. Comparisons were conducted between patients with and without immunoparesis, following the standard criteria for immunoparesis assessment^31^. The default MultiNicheNet pipeline (https://github.com/saeyslab/multinichenetr) was applied. To enhance robustness and prevent cluster drop-outs due to low numbers, certain subclusters were merged to form larger clusters per sample. Specifically, *“CD8 Effector_1*” and *“CD8 Effector_2*” were merged into the *“CD8_Effector cluster”*, “*CD8 Memory_1*” and “*CD8 Memory_2*” were merged into “*CD8_Memory cluster*”, *“cDC2*”and *“prog_DC*” were merged into “*DC cluster*”, “*prog_B 1*” and *“prog_B 2”* were merged into “*prog_B cluster*”, and “HSC”, “*LMPP*” and *“GMP*” were merged into “*HSC cluster*”, *“CD14 Mono*” and “*CD16 Mono*” were merged into “*monocyte cluster*”, and “*CD56 bright NK*” and “*NK*” were merged into “*NK cluster*”.

Clusters were included in the analysis only if a minimum of 4 cells were present in all samples per condition. gdT cells did not passed this filter, therefore not included in the Multinichenet analysis. Parameters were chosen carefully: LogFC_threshold was set to “0.5”, empirical_pval was set to “TRUE”, p_val_threshold was set to “0.05”, p_val_adj was set to “TRUE”, top_n_target was set to “250”, and prioritization weights were set to default.

All cell-cell interaction analyses and plots were conducted between clonal and polyclonal PC versus the entire TME. Clonal and polyclonal PCs were prioritized as sender cells, and the entire TME as receiver cells, and vice versa.

Top 75 ligands per condition were chosen for circos plots visualization and prioritized interactions for ligand-receptors pseudobulk expression products plots and ligand activity plots.

### Data visualization

All plots were generated using the ggplot2 (v 3.4.4) and ComplexHeatmap (v 2.18.0) packages in R.

Details on statistical tests used in the different figures and definition of relevant summary statistics are included in the figure legends.

## Supplementary figure legends

**Figure S1-S4. PCs heterogeneity elucidated by inferCNV**

InferCNV heatmap demonstrating of clonal (red) and polyclonal (blue) cells of each sample (n=46).

**Figure S5. GPRC5D and TNFRSF17 show differential expression in clonal and polyclonal cells**

**(A, B)** Violin plot of GPRC5D and TNFRSF17 gene expression pattern across disease stages (blue: HDs, green: MGUS, orange: SMM and red: MM)

**(C, D)** Violin plot of GPRC5D and TNFRSF17 gene expression pattern across disease stages (blue: HDs, green: MGUS, orange: SMM and red: MM) in clonal (red) and polyclonal (blue) cells.

**Figure S6. Quality control and pre-processing of single-cell polyclonal data**

**(A)** Violin plots illustrate distribution of percentage “nCounts_RNA” of each sample divided by HD, MGUS, SMM and MM

**(B)** Violin plots representing the distribution of mitochondrial genes percentage per each sample divided by HD, MGUS, SMM and MM

**(C)** Violin plots representing “nFeature_RNA” distribution of each sample divided by HD, MGUS, SMM and MM

**Figure S7. Polyclonal plasmacells clustering**

**(A, B)** Uniform Manifold Approximation and Projection (UMAP) representation of cells from the 64 single-cell RNA-seq data, colored by clusters **(A)** and by patients **(B).**

**Figure S8. Hallmark analysis of polyclonal plasmacells**

**(A, B, C)** Differential expression analysis showing Wilcoxon enriched hallmarks between HD *vs* MGUS **(A),** HD *vs* SMM **(B)** and HD *vs* MM **(C)** conditions. Gene-ratio on the y-axis, counts as dot size and color represent *padj* values in z-score.

**Figure S9. Cell abundance of patients’ plasma cells between immunoparesis conditions**

**(A, B)** Box plot representing patients’ clonal **(A)** and polyclonal **(B)** plasmacells distribution across immunoparesis negative (light-blue) or positive (yellow) patients.

**(C, D)** Donut plot showing the proportion of immunoparesis negative (light-blue) or positive (yellow) in patients’ clonal **(C)** and polyclonal (D) plasmacells and their respective number of cells.

**Figure S10. TME dataset integration and relative cell abundance of BM cell populations**

**(A)** UMAP of integrated CD138-cell fractions of n=31 samples colored by cell types.

**(B)** The box plot displays the relative abundance of cells within the integrated dataset (TME and PCs). Cell types are shown on the x-axis, with relative frequency represented on the y-axis in logarithm. The yellow boxes represent positive immunoparesis condition, while the lightblue boxes represent negative immunoparesis condition. The asterisk (*) above the “pPCs” label indicates a padj value of 0.004, determined by the Wilcoxon test with Benjamini-Hochberg adjustment.

**Figure S11. Scaled L-R pseudobulk products and ligand activity in PCs in immunoparesis-negative condition**

**(A, B)** Bubble plots illustrate scaled pseudobulk expression of ligand-receptor pairs specific for the “immunoparesis negative” condition. Rows represent ligand-receptor interactions, with sender and receiver cell types indicated respectively. PCs were set as senders in panel **A** and as receivers in panel **B.** Scaled ligand-receptor pseudobulk expression is depicted in the legend. The size of dots indicates whether a sample contained enough cells (>= 4) to be included in the differential expression (DE) analysis. On the right side, scaled ligand activity values, calculated based on the DE genes of cell type, and represented as normalized z-scores, are computed per receiver cell type. Higher values indicate greater enrichment of target genes of a specific ligand among the set of up-or downregulated genes in the “immunoparesis negative” or “positive” group. Gray color is associated to no DE genes (up or down) for those cell types due to pipeline parameters (see STAR methods).

**Figure S12. Predicted ligand-target interaction and average expression levels of target genes in pPCs of immunoparesis-negative patients**

**(A)** Heatmap showing predicted ligand-target links and the target regulatory potential behind ligand activity predictions. The genes shown are the top target genes (columns) of the ligand (rows) that have contributed to the ligand activity prediction of that interaction.

**(B)** Dot plot representing the average expression level of a target gene either up or down regulated in the predicted ligand-target plot. Upregulated genes are shown in red, while downregulated genes are shown in blue. Genes derive from pPCs L-R-T activity analyses in immunoparesis-negative condition.

**Figure S13. Scaled L-R pseudobulk products and ligand activity in PCs in immunoparesis-positive condition**

**(A,B)** Bubble plots illustrate scaled pseudobulk expression of ligand-receptor pairs specific for the “immunoparesis positive” condition. Rows represent ligand-receptor interactions, with sender and receiver cell types indicated respectively. PCs were set as senders in panel **A** and as receivers in panel **B**. Scaled ligand-receptor pseudobulk expression is depicted in the legend. The size of dots indicates whether a sample contained enough cells (>= 4) to be included in the differential expression (DE) analysis. On the right side, scaled ligand activity values, calculated based on the DE genes of cell type, and represented as normalized z-scores, are computed per receiver cell type. Higher values indicate greater enrichment of target genes of a specific ligand among the set of up-or downregulated genes in the “immunoparesis negative” or “positive” group. Gray color is associated to no DE genes (up or down) for those cell types due to pipeline parameters (see STAR methods).

**Figure S14. Predicted ligand-target link and average expression levels of target genes in pPCs of immunoparesis-positive patients**

**(A)** Heatmap showing predicted ligand-target links and the target regulatory potential behind ligand activity predictions. The genes shown are the top target genes (columns) of the ligand (rows) that have contributed to the ligand activity prediction of that interaction.

**(B)** Dot plot representing the average expression level of a target gene either up or down regulated regulated in the predicted ligand-target plot. Upregulated genes are shown in red, while downregulated genes are shown in blue. Genes derive from pPCs L-R-T activity analyses in immunoparesis-positive condition

