## Supplementary Table 1 for "Genotypic identification of polyclonal plasma cells in plasma cell dyscrasias shows an aberrant single-cell phenotype with clinical implications"

**Table S1: Patient characteristics**

| | MGUS | SMM | Active MM |
|---|---|---|---|
| **N. of patients** | 7 | 16 | 23 |
| **Female gender, n. (%)** | 1 (14) | 6 (38) | 10 (43) |
| **Age at sampling, median (range)** | 61 (41-83) | 65 (41-82) | 64 (48-84) |
| **Isotype, n. (%)** | | | |
| IgG-k | 3 (43) | 7 (44) | 12 (52) |
| IgA-k | 2 (29) | 3 (19) | 5 (22) |
| IgG-λ | 1 (14) | 5 (31) | 3 (13) |
| IgA-λ | 0 | 1 (6) | 1 (4) |
| Free light chain λ | 0 | 0 | 2 (9) |
| Free light chain k | 1 (14) | 0 | 0 |
| **Mayo Clinic Score, n. (%)** | | | |
| Low risk | 0 | - | - |
| Low-intermediate | 4 (57) | - | - |
| Intermediate-high | 3 (43) | - | - |
| High | 0 | - | - |
| **2-20-20 Score, n. (%)** | | | |
| Low | - | 6 (37.5) | - |
| Intermediate | - | 8 (50) | - |
| High | - | 2 (12.5) | - |
| **International Staging System, n. (%)** | | | |
| I | - | - | 13 (57) |
| II | - | - | 6 (26) |
| III | - | - | 4 (17) |
| **Revised International Staging System, n. (%)** | | | |
| I | - | - | 8 (34) |
| II | - | - | 11 (48) |
| III | - | - | 2 (9) |
| Not available, n (%) | - | - | 2 (9) |
| **IgH translocations, n. (%)** | | | |
| t(4;14) | 0 | 1 (6) | 2 (9) |
| t(11;14) | 1 (14) | 1 (6) | 4 (17) |
| t(14;16) | 1 (14) | 0 | 1 (4) |
| **Copy number abnormalities (FISH), n. (%)** | | | |
| gain(1q) | 0 | 0 | 3 (13) |
| amp(1q) | 0 | 3 (19) | 4 (17) |
| del(17p) | 1 (14) | 0 | 3 (13) |
| **FISH not available, n. (%)** | 3 (43) | 5 (31) | 2 (9) |
| **Immunoparesis, n. (%)** | | | |
| Involving ≥1 Ig class | 3 (43) | 14 (87.5) | 21 (91) |
