## Supplementary Table 3 for "Genotypic identification of polyclonal plasma cells in plasma cell dyscrasias shows an aberrant single-cell phenotype with clinical implications"

TABLES3

| celltype_pts | | | | | | | | | | | | | | | | | | | | | |
|---|---|---|---|---|---|---|---|---|---|---|---|---|---|---|---|---|---|---|---|---|---|
| | Immunoparesis | CD4 Memory | CD4 Naive | CD8 Naive | CD8_Effector | CD8_Memory | Clonal_Plasmablast | DC | gdT | HSC | MAIT | Memory B | Monocyte | Naive B | NK | pDC | Polyclonal_Plasmablast | Prog_B | Prog_Mk | Prog_RBC | Treg |
| **hr_mgus17** | NEG | 288 | 84 | 17 | 24 | 54 | 768 | 12 | 1 | 85 | 1 | 9 | 428 | 13 | 152 | 12 | 52 | 36 | 13 | 5 | 39 |
| **hr_mgus18** | NEG | 104 | 69 | 5 | 56 | 50 | 5541 | 4 | 0 | 149 | 4 | 5 | 213 | 9 | 104 | 16 | 236 | 8 | 8 | 10 | 12 |
| **hr_mgus24** | NEG | 249 | 33 | 37 | 45 | 94 | 1785 | 2 | 2 | 51 | 14 | 0 | 207 | 0 | 120 | 3 | 1262 | 15 | 0 | 23 | 6 |
| **hr_mgus4** | NEG | 222 | 124 | 12 | 101 | 187 | 1045 | 24 | 0 | 71 | 49 | 3 | 168 | 0 | 124 | 12 | 405 | 35 | 5 | 16 | 9 |
| **hr_mgus7** | NEG | 166 | 11 | 50 | 79 | 76 | 404 | 1 | 0 | 15 | 16 | 0 | 195 | 0 | 66 | 4 | 355 | 4 | 0 | 0 | 2 |
| **lr_mgus/mgrs16** | NEG | 29 | 29 | 1 | 16 | 50 | 1586 | 7 | 0 | 56 | 3 | 0 | 121 | 0 | 8 | 10 | 316 | 2 | 5 | 1 | 8 |
| **lr_mgus19** | NEG | 275 | 85 | 1 | 137 | 290 | 1650 | 22 | 1 | 110 | 9 | 12 | 504 | 18 | 15 | 19 | 477 | 28 | 13 | 23 | 9 |
| **mm10** | POS | 847 | 724 | 323 | 292 | 272 | 3818 | 21 | 21 | 88 | 58 | 60 | 171 | 159 | 784 | 123 | 21 | 97 | 16 | 26 | 6 |
| **mm12** | POS | 337 | 290 | 80 | 309 | 131 | 4141 | 3 | 4 | 35 | 35 | 51 | 124 | 51 | 19 | 0 | 23 | 1 | 8 | 6 | 0 |
| **mm19** | POS | 1121 | 318 | 191 | 321 | 375 | 2966 | 47 | 52 | 355 | 71 | 2 | 370 | 27 | 463 | 26 | 53 | 69 | 27 | 47 | 14 |
| **mm23** | POS | 1071 | 189 | 83 | 681 | 612 | 2584 | 16 | 3 | 135 | 143 | 1263 | 782 | 254 | 461 | 11 | 24 | 13 | 20 | 5 | 16 |
| **mm24** | POS | 165 | 18 | 1 | 422 | 921 | 6624 | 7 | 0 | 4 | 2 | 29 | 28 | 0 | 273 | 0 | 52 | 0 | 6 | 2 | 6 |
| **mm27** | POS | 2231 | 52 | 20 | 83 | 738 | 3930 | 6 | 27 | 66 | 631 | 41 | 1327 | 27 | 1510 | 40 | 10 | 0 | 48 | 91 | 33 |
| **mm9** | POS | 1137 | 299 | 45 | 476 | 124 | 2185 | 32 | 5 | 60 | 79 | 16 | 189 | 1 | 204 | 75 | 53 | 3 | 21 | 25 | 58 |
| **smm10_progression** | POS | 742 | 58 | 2 | 283 | 645 | 5043 | 9 | 0 | 117 | 56 | 3 | 1975 | 18 | 215 | 24 | 1562 | 38 | 16 | 33 | 28 |
| **smm100** | POS | 223 | 35 | 10 | 89 | 184 | 5216 | 32 | 2 | 363 | 25 | 3 | 791 | 0 | 51 | 8 | 225 | 0 | 3 | 14 | 30 |
| **smm11** | POS | 592 | 80 | 119 | 52 | 82 | 406 | 3 | 9 | 29 | 92 | 1 | 254 | 2 | 333 | 6 | 65 | 4 | 2 | 5 | 4 |
| **smm12** | POS | 226 | 81 | 7 | 227 | 105 | 4908 | 9 | 8 | 74 | 41 | 123 | 401 | 22 | 148 | 7 | 261 | 7 | 10 | 7 | 9 |
| **smm14** | POS | 89 | 11 | 6 | 118 | 25 | 3275 | 9 | 2 | 122 | 33 | 0 | 319 | 0 | 79 | 16 | 57 | 23 | 8 | 23 | 15 |
| **smm2** | POS | 221 | 19 | 0 | 149 | 440 | 4354 | 10 | 2 | 38 | 10 | 6 | 319 | 0 | 51 | 16 | 97 | 20 | 12 | 8 | 8 |
| **smm2_progression** | POS | 111 | 23 | 9 | 29 | 177 | 6403 | 8 | 3 | 25 | 13 | 5 | 247 | 0 | 26 | 3 | 146 | 1 | 43 | 40 | 6 |
| **smm20** | POS | 448 | 106 | 0 | 115 | 231 | 3597 | 15 | 4 | 148 | 13 | 3 | 906 | 4 | 113 | 18 | 210 | 2 | 13 | 10 | 32 |
| **smm21** | POS | 247 | 126 | 25 | 31 | 347 | 637 | 30 | 1 | 79 | 2 | 2 | 293 | 2 | 126 | 16 | 201 | 21 | 15 | 28 | 9 |
| **smm23** | POS | 448 | 96 | 21 | 155 | 466 | 3860 | 11 | 1 | 97 | 13 | 1 | 609 | 2 | 230 | 10 | 190 | 23 | 5 | 15 | 5 |
| **smm28** | POS | 262 | 142 | 8 | 28 | 524 | 4011 | 0 | 7 | 38 | 18 | 1 | 895 | 3 | 148 | 15 | 66 | 5 | 16 | 39 | 23 |
| **smm3** | POS | 88 | 34 | 2 | 15 | 15 | 2533 | 14 | 3 | 43 | 1 | 1 | 208 | 9 | 138 | 12 | 52 | 29 | 4 | 1 | 4 |
| **smm3_interim** | POS | 927 | 473 | 101 | 302 | 119 | 7683 | 61 | 8 | 319 | 55 | 27 | 650 | 363 | 461 | 31 | 62 | 218 | 15 | 47 | 27 |
| **smm5** | NEG | 105 | 44 | 8 | 27 | 25 | 378 | 4 | 0 | 25 | 42 | 0 | 127 | 0 | 20 | 10 | 421 | 21 | 6 | 11 | 2 |
| **smm6** | POS | 102 | 10 | 1 | 14 | 47 | 1962 | 3 | 0 | 55 | 4 | 1 | 87 | 8 | 62 | 6 | 184 | 28 | 2 | 9 | 3 |
| **smm6_progression** | POS | 215 | 6 | 0 | 52 | 56 | 2681 | 1 | 0 | 43 | 13 | 6 | 211 | 4 | 58 | 3 | 237 | 31 | 13 | 13 | 2 |
| **smm9** | POS | 236 | 64 | 5 | 77 | 99 | 6220 | 22 | 1 | 85 | 2 | 0 | 279 | 6 | 27 | 22 | 72 | 14 | 13 | 894 | 13 |
