## Supplementary Table S4 for "Genotypic identification of polyclonal plasma cells in plasma cell dyscrasias shows an aberrant single-cell phenotype with clinical implications"

TABLES4

| | |
|---|---|
| 1 | BCL2L11 |
| 2 | CCDC85B |
| 3 | COBLL1 |
| 4 | DNAAF1 |
| 5 | EIF2AK3 |
| 6 | FNBP1 |
| 7 | HCST |
| 8 | HIST1H1C |
| 9 | JSRP1 |
| 10 | KHDRBS1 |
| 11 | NEAT1 |
| 12 | RHOH |
| 13 | RPL10 |
| 14 | RPL10A |
| 15 | RPL15 |
| 16 | RPL21 |
| 17 | RPLP2 |
| 18 | RPS16 |
| 19 | RPS24 |
| 20 | UBALD2 |
| 21 | UBE2J1 |
| 22 | UQCRB |
| 23 | ACADM |
| 24 | ANXA6 |
| 25 | ATM |
| 26 | AURKAIP1 |
| 27 | CALM3 |
| 28 | COA3 |
| 29 | COX7B |
| 30 | CPNE5 |
| 31 | CTSH |
| 32 | CYC1 |

| | |
|---|---|
| 33 | DNAJC1 |
| 34 | DNAJC19 |
| 35 | DUSP22 |
| 36 | EIF2AK1 |
| 37 | FOXO3 |
| 38 | GADD45GIP1 |
| 39 | GFER |
| 40 | GMPPA |
| 41 | HSPB11 |
| 42 | IFI27L2 |
| 43 | IKBIP |
| 44 | IRF2 |
| 45 | ISOC2 |
| 46 | KLHL5 |
| 47 | MANEA |
| 48 | MDH1 |
| 49 | METTL23 |
| 50 | MPHOSPH8 |
| 51 | MRPL20 |
| 52 | MRPS34 |
| 53 | MRPS36 |
| 54 | NDUFA8 |
| 55 | NDUFB2 |
| 56 | NDUFB4 |
| 57 | NDUFV2 |
| 58 | OST4 |
| 59 | PARP1 |
| 60 | PLEK |
| 61 | PNOC |
| 62 | POLR2F |
| 63 | PPCDC |
| 64 | PPM1K |
| 65 | PRDX3 |
| 66 | PRKAR1A |

| | |
|---|---|
| 67 | PRR34-AS1 |
| 68 | PSMA4 |
| 69 | PSMA5 |
| 70 | PSMB9 |
| 71 | PTPN6 |
| 72 | PYCARD |
| 73 | RPL22L1 |
| 74 | RSRP1 |
| 75 | SAR1B |
| 76 | SERF2 |
| 77 | SMCHD1 |
| 78 | SPATS2 |
| 79 | SYTL1 |
| 80 | TIMM13 |
| 81 | TXNDC15 |
| 82 | UCP2 |
| 83 | UQCC2 |
| 84 | UQCRQ |
| 85 | ZBP1 |
| 86 | ZBTB8OS |
| 87 | ZNF706 |
| 88 | ZNHIT1 |
| 89 | ZRANB2 |
| 90 | AC103591.3 |
| 91 | AC233755.2 |
| 92 | AMPD1 |
| 93 | AP1S2 |
| 95 | ARMC10 |
| 96 | ASNSD1 |
| 97 | ATXN10 |
| 98 | BNIP3 |
| 99 | BRK1 |
| 100 | C15orf61 |
| 101 | CADPS2 |

| | |
|---|---|
| 102 | CAMTA1 |
| 103 | CCDC88C |
| 104 | CMTM7 |
| 105 | CSRNP1 |
| 106 | CTHRC1 |
| 107 | CTSW |
| 108 | DDT |
| 109 | DNAJB11 |
| 110 | DSTN |
| 111 | EIF4G2 |
| 112 | GLO1 |
| 113 | GLS |
| 114 | HMGN2 |
| 115 | HSP90B1 |
| 116 | IFT27 |
| 117 | ITGA6 |
| 118 | ITGA8 |
| 119 | LYZ |
| 120 | MOXD1 |
| 121 | MRPL33 |
| 122 | NDUFA5 |
| 123 | NME3 |
| 124 | NUS1 |
| 125 | PARVB |
| 126 | PDIA4 |
| 127 | PGRMC2 |
| 128 | PHF14 |
| 129 | PLA2G2D |
| 130 | PSMA3-AS1 |
| 131 | PTK2B |
| 132 | RAB11FIP1 |
| 133 | RBBP6 |
| 134 | SEC22B |
| 135 | SEC24D |

| | |
|---|---|
| 136 | SEC61G |
| 137 | SF3B5 |
| 138 | SNHG9 |
| 139 | SOD1 |
| 140 | SPG21 |
| 141 | SRP9 |
| 142 | SRSF5 |
| 143 | SRSF7 |
| 144 | TMED2 |
| 145 | TMEM50B |
| 146 | TRA2A |
| 147 | UFM1 |
| 148 | Z93241.1 |
| 149 | ZFP36 |
